# Genetic architecture underlying changes in carotenoid accumulation during the evolution of the Blind Mexican cavefish, *Astyanax mexicanus*

**DOI:** 10.1101/788844

**Authors:** Misty R. Riddle, Ariel Aspiras, Fleur Damen, John N. Hutchinson, Daniel Chinnapen, Clifford J. Tabin

**Author notes:** Department of Stem Cell and Regenerative Biology, Harvard University, Cambridge, MA 02138.

## Abstract

Carotenoids are yellow to orange pigments produced by plants, bacteria, and fungi. They are consumed by animals and metabolized to produce molecules essential for gene regulation, vision, and pigmentation. Cave animals represent an interesting opportunity to understand how carotenoid utilization evolves. Caves are devoid of light, eliminating primary production of energy through photosynthesis and therefore limiting carotenoid availability. Moreover, the selective pressures that favor carotenoid-based traits, like pigmentation and vision, are relaxed. *Astyanax mexicanus* is a species of fish with river-adapted (surface) and multiple cave-adapted populations (i.e. Tinaja, Pachón, Molino). Cavefish exhibit regressive features such as loss of eyes and melanin pigment, and constructive traits, like increased sensory neuromasts and starvation resistance. Here we show that unlike surface fish, Tinaja and Pachón cavefish accumulate carotenoids in the visceral adipose tissue (VAT). Carotenoid accumulation is not observed in Molino cavefish indicating that it is not an obligatory consequence of eye loss. We used quantitative trait loci mapping and RNA sequencing to investigate genetic changes associated with this trait. Our findings suggest that multiple stages of carotenoid processing may be altered in cavefish, including absorption and transport of lipids, cleavage of carotenoids into un-pigmented molecules, and differential development of intestinal cell types involved in carotenoid assimilation. Our study establishes *A. mexicanus* as a model to study the genetic basis of natural variation in carotenoid accumulation and how it impacts physiology.

**Research Highlights:**

- Cavefish accumulate carotenoids in the visceral adipose tissue (VAT)
- Genetic mapping reveals candidate genes associated with yellow VAT
- Carotenoid accumulation is linked with decreased expression of carotenoid-processing genes

## Introduction

Carotenoids are an important class of molecules with conjugated double bonds that absorb visible light and appear yellow to red. They are synthesized by plants and consumed by animals either directly from eating plants, or from consuming plant-eating animals. Carotenoid metabolism produces the chromophore for all known visual systems (11-cis-retinal, Vitamin A) and an essential regulator of development and gene expression (retinoic acid). Differential processing and distribution of carotenoids is responsible for the spectacular range of yellow to red colors observed in bird plumage, fish pigment, and reptile skin (reviewed in Toews et al., 2017). In humans, carotenoid levels have been linked to risk for certain cancers, cardiovascular disease, and macular degeneration (reviewed in Eggersdorfer and Wyss, 2018). Despite interest in understanding natural variation in carotenoid processing and utilization, there are a limited number of genetic models.

Carotenoid-based traits are influenced by many variables. Animal diets differ in carotenoid abundance over space and time making it difficult to make functional comparisons in wild populations. Moreover, how carotenoids are digested, absorbed, processed, and translocated likely involves many genes. Once carotenoids are released from the food matrix, they are mixed with bile salts and fats to form micelles that are absorbed into enterocytes by the action of lipid transporters. Carotenoids can be cleaved within the enterocyte or packaged into chylomicrons that are transferred into the lymphatic system. Chylomicrons are differentially routed to tissues based on tissue-specific expression of lipoprotein receptors. Genetic variation at each of these steps has been linked to changes in carotenoid-based traits (reviewed in Moran et al., 2018; Toews et al., 2017). For example, mutations in β-carotene oxygenase (BCO1) and β-carotene-9′,10′-dioxygenase (BCO2), enzymes that convert carotenoids into unpigmented metabolites, are linked with yellow phenotypes in chickens, sheep, and cows (Berry et al., 2009; Eriksson et al., 2008; Våge & Boman, 2010). Mutations in the lipid transporter SCARB1 produce pale-colored silk-worms due to decreased absorption of β-carotene (Sakudoh et al., 2013) and high expression of SCARB1 in the intestine of salmon is associated with increased carotenoid accumulation and orange muscle color (Sundvold, Helgeland, Baranski, Omholt, & Våge, 2011). In humans, single nucleotide polymorphisms in BCO1, BCO2, SCARB1 and other lipid transporters and are associated with carotenoid levels in the serum (reviewed in Moran et al., 2018). Considering the potential health benefits of carotenoids, there is commercial interest in breeding animals that produce high carotenoid levels as well as understanding what causes variation in human carotenoid absorption.

Birds, reptiles and fish utilize carotenoids to produce visual signals that are used to avoid predators and attract mates (Toews et al., 2017). Extensive research on the elaboration of carotenoid-based pigment in cichlid fish has revealed associations between carotenoid processing and mate-preference, social dominance, and oxidative stress (reviewed in Sefc et al., 2014). A combination of genetic mapping and gene expression studies in cichlids indicate that changes in genes that influence carotenoid metabolism (*bco2*), or the development of pigment-containing xanthaphore cells (*csf1ra*, *Edn3b*, *Pax7*) produce variation in carotenoid-based pigment. In stickleback fish, throat and pelvic fin carotenoid coloration genetically maps to three regions on the genome (Yong, Peichel, & McKinnon, 2016). The evolutionary trade-off of carotenoid-coloration with other phenotypes dependent on carotenoids like vision and anti-oxidant function is difficult to dissect.

Studying carotenoid utilization in cave species offers a novel opportunity to disentangle the variables that contribute to carotenoid-based traits. Cave environments are devoid of light, eliminating the favorability of pigmentation and vision but also limiting carotenoid availability. Wolfe and Cornwell (1964) compared carotenoid levels in cave-adapted and non-cave-adapted species of crayfish found in the same cave in Kentucky. They found two pigments in the cave form (β-carotene and lutein) and eight in the surface form. The level of β-carotene was over six times higher in the cave-adapted form and the level of lutein was over twelve times higher. The authors concluded that the cave forms lost the ability to generate keto-carotenoids from dietary carotenoids but the genetic basis of this difference and impact on other crayfish phenotypes was not investigated.

The Mexican tetra, *Astyanax mexicanus* is a powerful model to study the genetic basis of evolution. It is a single species of fish that exists as a river-adapted form (surface fish) found in rivers from Central Mexico to Southern Texas, and a cave-adapted form (cavefish) found in perpetually dark limestone caves of the Sierra del Abra region in Northeastern Mexico (Elliott, 2015). Surface fish and cavefish have adapted to dramatically different diets; river fish consume plants and insects throughout the year, while most cavefish populations are dependent on external sources of food brought in by seasonal flooding or bat droppings (Turner, 2017). There are 29 known cave populations that are of polyphyletic origin. Of the three cave populations used in this study, those from the Tinaja and Pachón caves form a clade separate from Molino cavefish (Herman et al., 2018). The populations are interfertile and amenable to genetic manipulation (Stahl et al., 2019) allowing researchers to investigate the genetic changes associated with behavioral (Chin et al., 2018; Hyacinthe, Attia, & Rétaux, 2019; Lloyd et al., 2018; Yoshizawa et al., 2015) metabolic (Aspiras, Rohner, Martineau, Borowsky, & Tabin, 2015; Riddle, Aspiras, et al., 2018; Xiong, Krishnan, Peuß, & Rohner, 2018), and morphological (Gross & Powers, 2018; Lyon, Powers, Gross, & O’Quin, 2017) cave-adapted traits, and utilize the different cave populations as natural replicates.

While understanding eye and pigment regression has been a major focus in *A. mexicanus* research (Gore et al., 2018; Jeffery, Ma, Parkhurst, & Bilandžija, 2016; Klaassen, Wang, Adamski, Rohner, & Kowalko, 2018; Krishnan & Rohner, 2017), there is a dearth of knowledge on carotenoid processing. It has been noted that cave populations exhibit increased deposits of yellow-orange color (Hinaux et al., 2011), but the molecular and genetic basis of this difference is unclear. Based on whole transcriptome comparison at multiple developmental stages, Pachón cavefish have greater expression of an enzyme that converts ß-carotene into colorless compounds (BCO2), a surprising finding considering Pachón larvae appear more yellow (Stahl & Gross, 2017). How carotenoid-processing evolved in cave-adapted forms of *A. mexicanus*, and its potential influence on other traits, has remained unexplored.

Here we present the first analysis of carotenoid processing in laboratory-raised *A. mexicanus*. We show that both surface fish and cavefish exhibit carotenoid-based pigment. We find that unlike surface fish, Pachón and Tinaja cavefish accumulate carotenoids in the visceral adipose tissue (VAT). We show that accumulation of carotenoids is not associated with eye loss or changes in the length of the digestive tract. Using genetic mapping, we discovered a quantitative trait locus associated with yellow fat and identified coding mutations in a gene that could influence absorption of carotenoids by altering the number of absorptive cell-types in the intestine. We show that Tinaja cavefish have reduced expression of genes important for carotenoid absorption, metabolism, and transport. Finally, we discuss how these differences may have evolved and the potential impacts on cavefish physiology.

## Results

Surface fish exhibit carotenoid-based pigmentation on the tail and anal fin (Figure 1A). Based upon laboratory observation, anal fin color may be associated with social interaction. When housed together in the laboratory, the surface fish that exhibits the most aggressive behavior develops a bright orange stripe on the anterior edge of the anal fin (Supplementary Video 1). Interestingly, if surface fish are housed individually they each develop a bright orange stripe (Figure 1A). Tinaja, Pachón, and Molino cavefish also exhibit orange pigmentation on the tail and anal fin, although it tends to not be as pronounced (Figure 1A, 3A). Quantifying carotenoid-based pigmentation in the integument is challenging due to the presence of melanophores and iridiphores in intervening layers. To investigate carotenoid processing in *A. mexicanus*, we therefore started by examining pigments in the internal organs.

**Figure 1.**
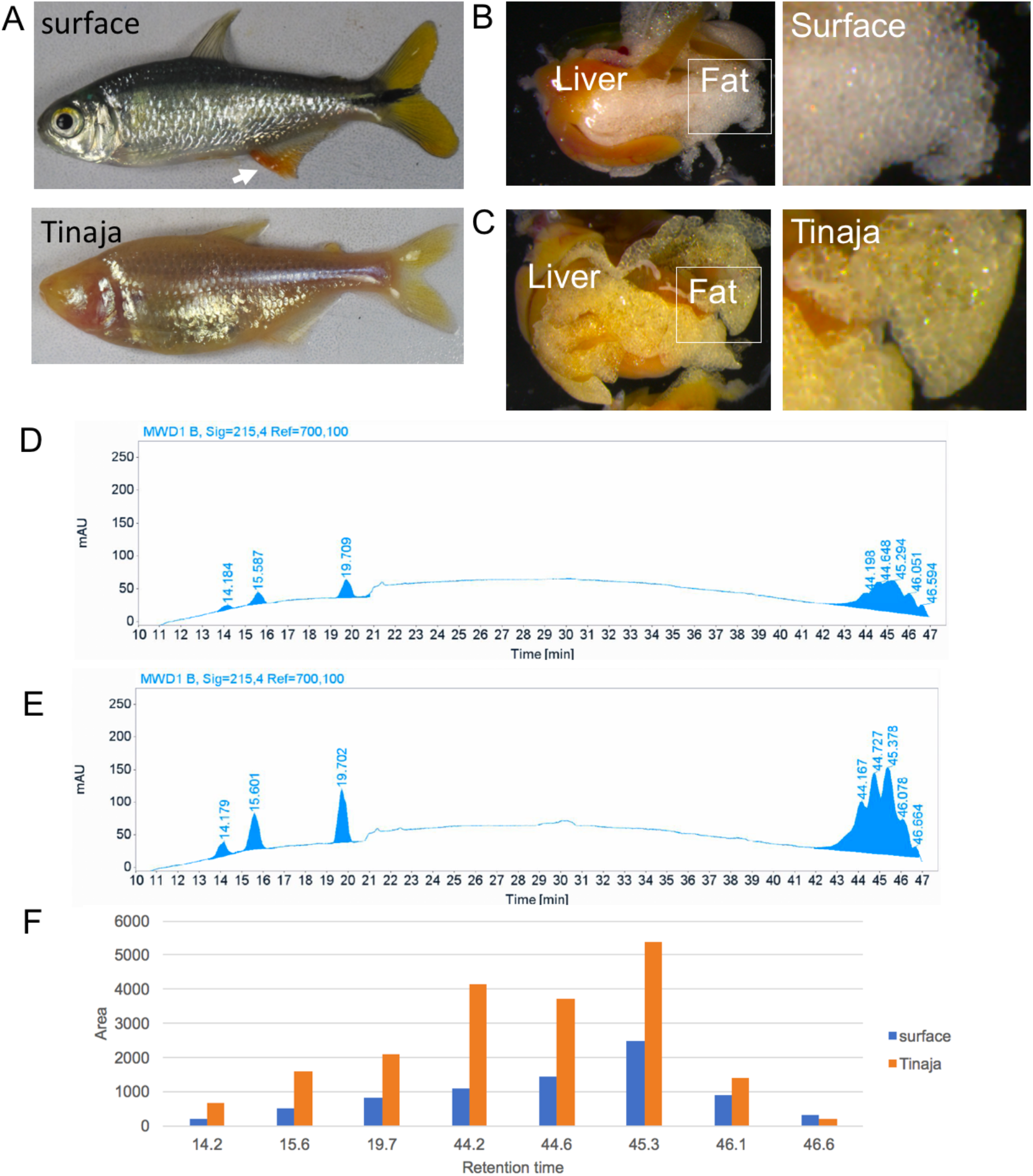
*Astyanax mexicanus* cavefish accumulate yellow pigment in the visceral adipose tissue (VAT). A, image of surface fish (top) and Tinaja cavefish (bottom). B-C, Visceral organs of surface fish and Tinaja cavefish (right panel shows larger image of VAT). D-E, HPLC analysis of carotenoids extracted from VAT of surface fish (D) and Tinaja cavefish (E). F, Bar graph showing comparing peak area at the indicated retention time.

### Tinaja cavefish accumulate yellow pigments in the visceral adipose tissue (VAT)

We dissected lab-raised surface fish and Tinaja cavefish that were fed *ad libitum* with a combination of brine shrimp and commercially available pellet food that is supplemented with β-carotene (see methods). We observed a striking difference in the color of the fat surrounding the internal organs; the VAT of the Tinaja cavefish is characterized by a markedly yellow color (Figure 1B, C). The yellow color led us to hypothesize that it represents an accumulation of dietary carotenoids. To test this hypothesis, we fed fish a diet that is comparatively low in carotenoids: small white non-parasitic worms *Enchytraeus buchholzi* (grindal worms). We found that Tinaja cavefish do not have yellow VAT on this diet (Figure 2).

**Figure 2.**
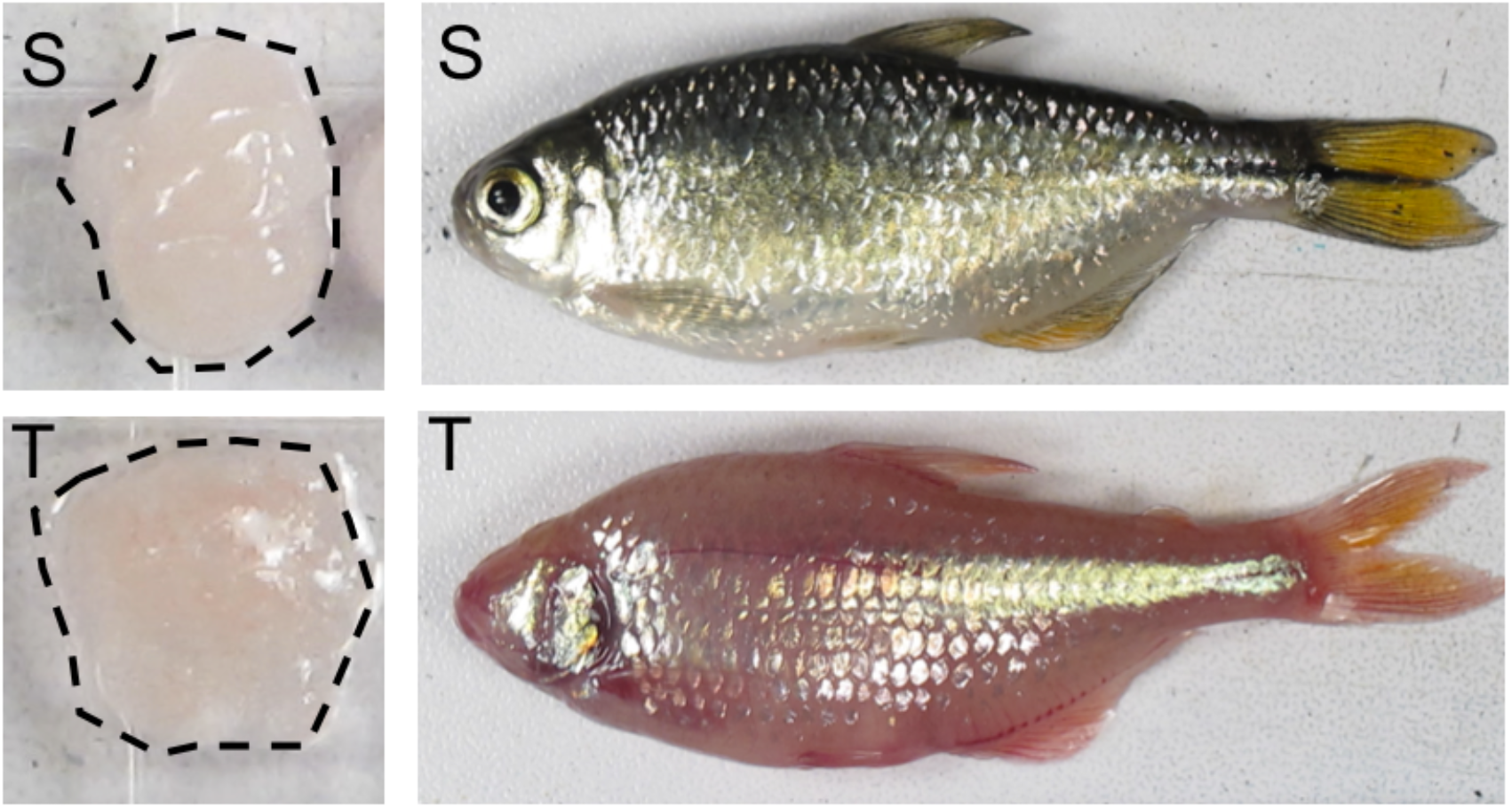
Color of visceral adipose tissue on low-carotenoid diet. Images of surface fish (S) and Tinaja cavefish (T) VAT (outlined with dotted black line). Fish fed a diet of white worms (*Enchytraeus buchholzi*).

We considered that similar to cave-adapted crayfish, Tinaja may have lost the ability to produce unpigmented molecules like retinol from carotenoids. We extracted pigments from the same amount of VAT from surface fish and Tinaja cavefish fed the high-carotenoid diet and analyzed the samples using High-pressure liquid chromatography (HPLC, see Methods). We determined the mobility of retinol and β-carotene as standards within the same experiment. We observed compounds with the same retention times in surface fish and Tinaja cavefish samples (Figure 1 D-E). Three peaks have retention times of less than 21 minutes (14.2, 15.6, 19.7), representing pigments with similar properties to retinol (elution time of 18.8 minutes) and five peaks have elution times of greater than 41 minutes (44.2, 44.7, 45.3, 46.1, 46.6), representing pigments with similar properties to β-carotene (elution time of 42.2). We found that the height and area of all but one peak (retention time 46.6) is greater in the Tinaja samples (Figure 1F). These results suggest that surface fish and cavefish have the same pigments in the VAT, but Tinaja may have a greater concentration.

### Variables associated with accumulation of carotenoids

We next investigated the variables that could influence carotenoid accumulation in the VAT. We first considered the impact of appetite. Tinaja cavefish evolved hyperphagia as an adaptation to the infrequent influx of nutrients experienced in the cave environment (Aspiras et al., 2015). To control for differences in food consumption, we individually housed surface fish and cavefish and ensured they consumed 6 mg of pellet food per day for greater than four months. While the color difference of the VAT was not as striking to fish fed *ad libitum*, we found that, most Tinaja had VAT that was visibly more yellow compared to surface fish (Supplemental Figure 1). These findings indicate that differences in appetite are not sufficient to explain carotenoid accumulation.

The main cleavage product of carotenoids, retinol, is essential for vision. We considered that lack of eyes may lead to an accumulation of metabolites that are going unused in eyeless cavefish. We examined VAT color in two additional cave populations that have complete eye regression and found that fish from the Pachón cave have yellow VAT similar to Tinaja, but Molino cavefish have white VAT (Figure 3A). To further dissect the interaction between eye degeneration and carotenoid accumulation we interbred surface fish and Tinaja cavefish to create F1 hybrids which have smaller eyes than surface fish. We found that one out of three hybrids had yellow VAT (Supplemental Figure 1). The hybrid with the yellow pigment also had a growth in the abdomen, introducing an additional variable that appears to influence VAT color. We interbred F1 hybrids and analyzed VAT color and eye diameter in the surface/Tinaja F2 hybrid population that were individual housed and consumed the same amount per day for greater than four months. In hybrids that had enough fat to analyze (124/217) we scored VAT color as yellow (n=12) and not yellow (n=112, Figure 3B). We found that some surface/cave F2 hybrids that lacked eyes had white fat, and vice versa, some fish with eyes had yellow fat (Figure 3B). In addition, we found that average relative eye diameter did not differ between fish with yellow or white VAT (left eye: 0.062 vs 0.065, p-value = 0.34, right eye: 0.066 vs 0.063, p-value = 0.35, Wilcox-rank test). These combined results suggest that eye loss is not sufficient to explain accumulation of carotenoids, substantiating the conclusion we reached based on the Molino cavefish.

**Figure 3.**
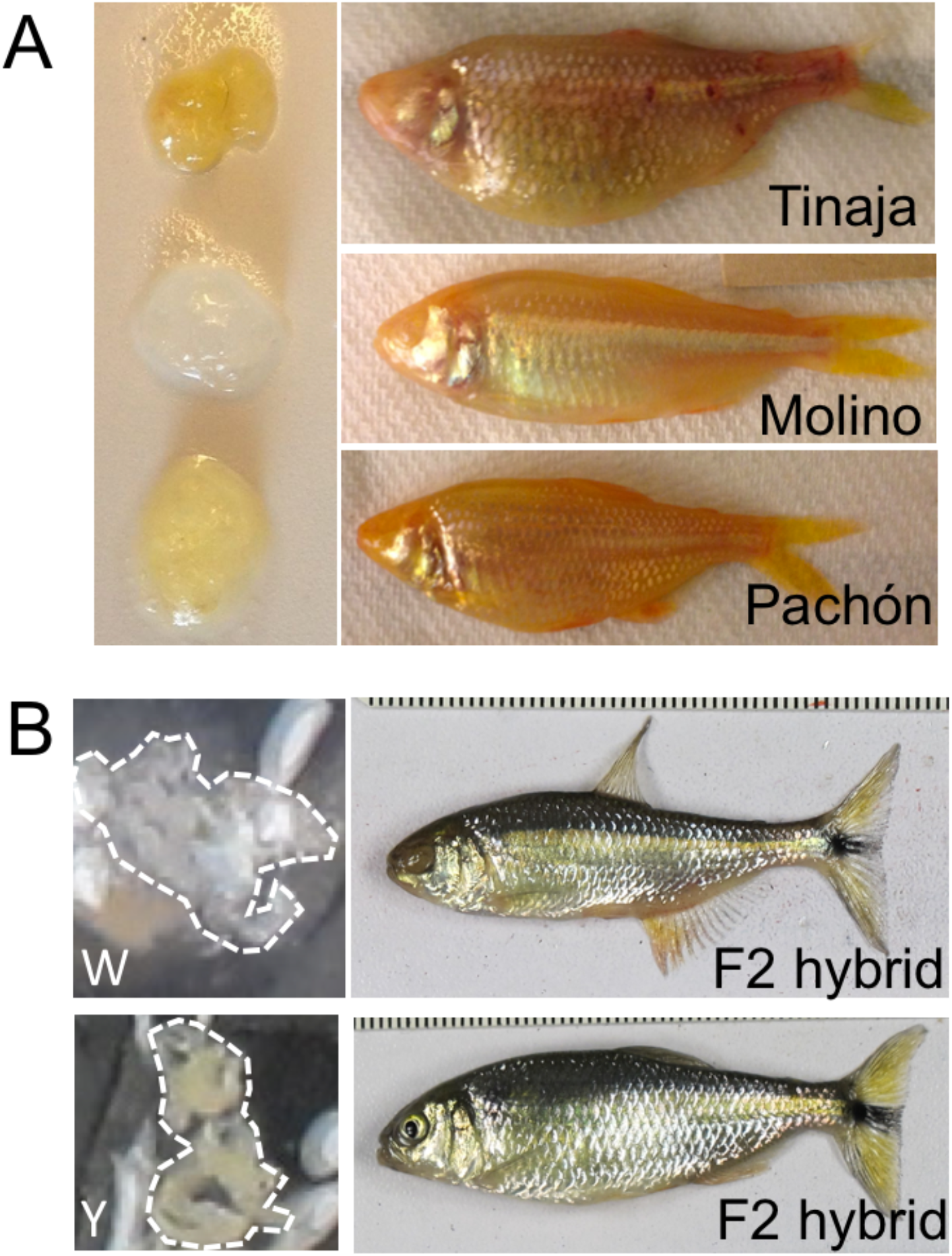
Eye degeneration is not associated with carotenoid accumulation in *A. mexicanus*. A, Images of VAT from individuals of the indicated cave populations. B, Images of VAT from eyeless (top) and eyed (bottom) fish from surface/Tinaja F2 hybrid population. VAT is outlined by white dotted line, W indicates the white sample taken from the eyeless F2 hybrid, and Y indicates the yellow sample taken from the eyed F2 hybrid.

We next considered differences in absorptive capacity of the gastrointestinal tract. Carotenoids are packaged into micelles that are absorbed by enterocytes in the intestine. The surface area available for absorption could, in principle, lead to a difference in carotenoid accumulation. To determine if VAT color and the length of the gastrointestinal tract are associated in *A. mexicanus* we examined these traits in the surface/Tinaja F2 hybrids. We measured the length of each portion of the gut (stomach, midgut and hindgut) from macroscopic images (Supplemental Figure 2) and normalized the values to fish length. We did not find a significant difference in the relative gut length of any region between hybrids with yellow fat versus white fat (relative total gut length 0.688, 0.679 respectively, p=0.78 Wilcox-rank test).

### Genetic mapping reveals quantitative trait loci associated with yellow VAT

To search for genes associated with carotenoid accumulation we carried out a quantitative trait loci (QTL) analysis using F2 surface/Tinaja hybrids (see methods). We scored fat color as a binary trait and using a single-QTL model identified a QTL on linkage group 16 with a LOD score of 8.0 at marker r164788 accounting for 25.53% of the variance (Figure 4). Five markers on linkage group 16 have LOD scores above the 5% significance threshold of 3.95 (Table 1). To identify candidate genes that contribute to fat color, we determined the position of the markers on the surface fish genome assembly. Two of the markers are on the same chromosome and three are on separate unplaced scaffolds. We identified the genes in the region between the markers on the same chromosome (3: 2805128-3751814). There are eight genes in this region: *clip2*, *gtf2ird1*, *DTX2*, *tmem132e*, and four that are uncharacterized (Table 2).

**Table 1.** QTL markers for yellow fat phenotype. Marker name and position on the linkage group (LG) and surface fish chromosome.

| marker | LOD | LG: position | Chromosome: position |
| --- | --- | --- | --- |
| r179950 | 7.8 | 16:20.1 | 3: 2805128 |
| r180290 | 5.8 | 16:23.5 | 3: 3751814 |
| r165238 | 3.4 | 16:01.4 | APWO02000155: 987504 |
| r164788 | 8.0 | 16:18.4 | APO020000139:6870990 |
| r9999 | 7.3 | 16:11.0 | APWO02000113: 510973 |

**Table 2.** Candidate genes for fat color QTL. Gene IDs, name, and available description of genes within QTL peak (chromosome 3: 2805128-3751814).

| Gene stable ID | Gene name | Description |
| --- | --- | --- |
| ENSAMXG00000007486 | DTX2 | E3 ubiquitin-protein ligase |
| ENSAMXG00000007478 | Clip-2 | CAP-Gly domain containing linker protein 2 |
| ENSAMXG00000007488 | tmem132e | Transmembrane protein |
| ENSAMXG00000007482 | GTF2I | GTF2I repeat domain containing 1 |
| ENSAMXG00000033138 |  | Retrotransposable element |
| ENSAMXG00000037591 |  | Transposase derived nuclease |
| ENSAMXG00000029351 |  |  |
| ENSAMXG00000038139 |  |  |

**Figure 4:**
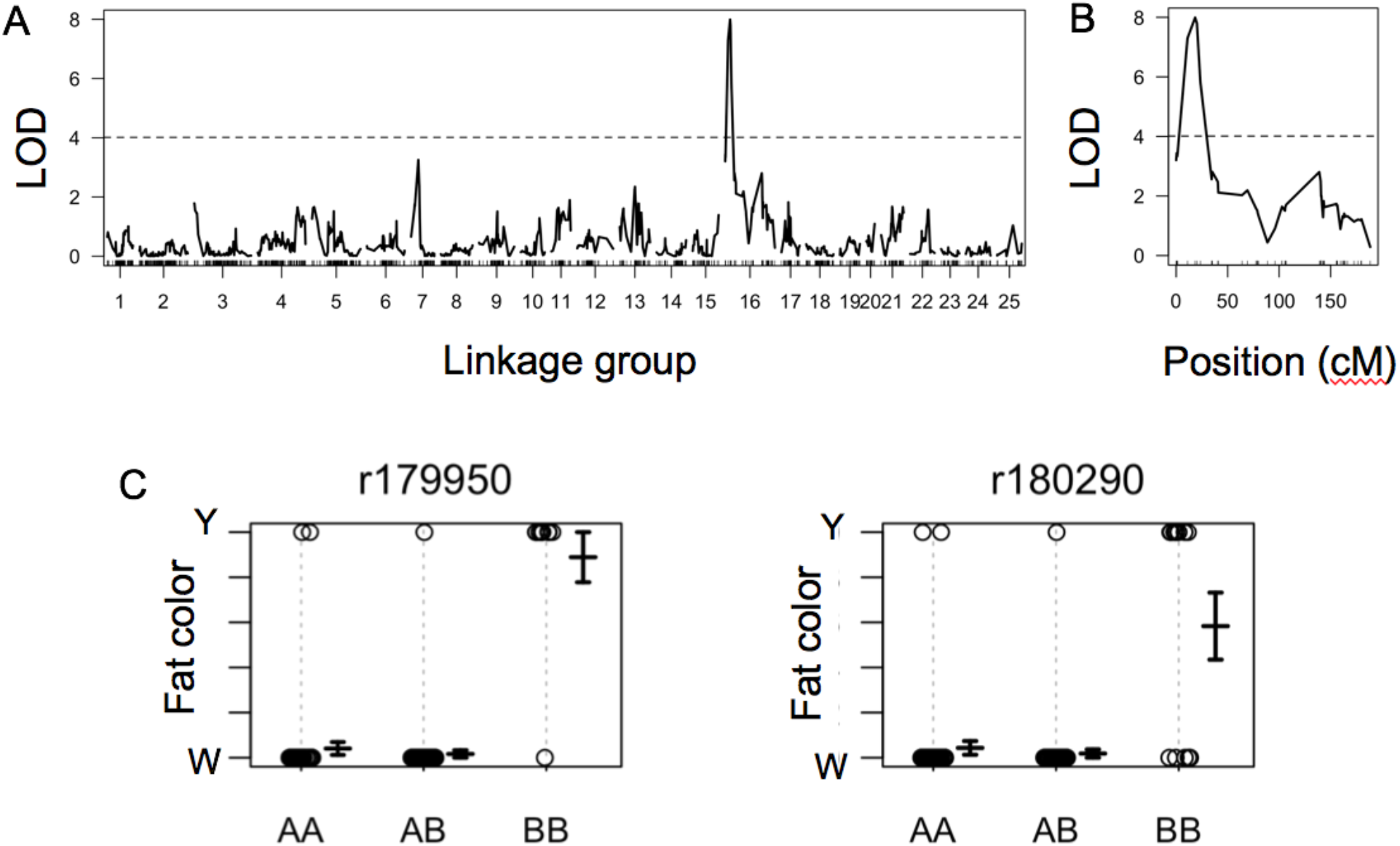
QTL associated with yellow fat color. A, Results of genome wide LOD calculation using Haley-Knott regression and binary model. B, LOD scores for markers on linkage group 16. Significance threshold of 5% (black dotted line) C, Genotype and phenotype at indicated maker positions (AA, surface; BB, cave).

We considered which of these genes could influence carotenoid processing based on known activities. *Clip-2* and *tmem132e* likely function in neurons and have roles in microtubule stability (Vandeweyer, Van Der Aa, Reyniers, & Kooy, 2012) and hair cell function respectively (Sanchez-Pulido & Ponting, 2018). GTF2I is a general transcription factor that is ubiquitously expressed. DTX2 is an E3 ubiquitin ligase that acts as a positive regulatory of Notch signaling (Matsuno, Diederich, Go, Blaumueller, & Artavanis-Tsakonas, 1995) and is expressed in the mouse and human gastrointestinal (GI) tract (Smith et al., 2019)(GTEx Analysis Release V8 (dbGaP Accession phs000424.v8.p2)). Considering that Notch activity promotes the formation of absorptive cells in the GI tract (Tian et al., 2015) we compared surface fish and Pachón cavefish DTX2 using the available annotations for the surface fish and Pachón cavefish genomes (Figure 5A). Surface fish have two splice isoforms of DTX2. We aligned the predicted protein sequences and found that the first 39 and last 12 amino acids are absent in Pachón cavefish. Pachón cavefish have a valine to isoleucine substitution (V166I) in the WWE protein domain that is important for Notch interaction (AA29-195). Fish from this population also have two substitutions R441H, S443L in the ring-finger domain (AA435-497) and an additional substitution P398A that is not in a specific domain. We found that Pachón cavefish also have additional amino acids that create a predicted transmembrane domain (Figure 5A,B)(Sonnhammer, von Heijne, & Krogh, 1998). One of the insertions represents a difference in the cDNA sequence and the addition of two exons. The other change is an addition of a repeated 20 base pair (bp) sequence in the genomic DNA (CTGCAAACTTTTCGTCCT); surface fish have two repeats and Pachón cavefish have three.

**Figure 5.**
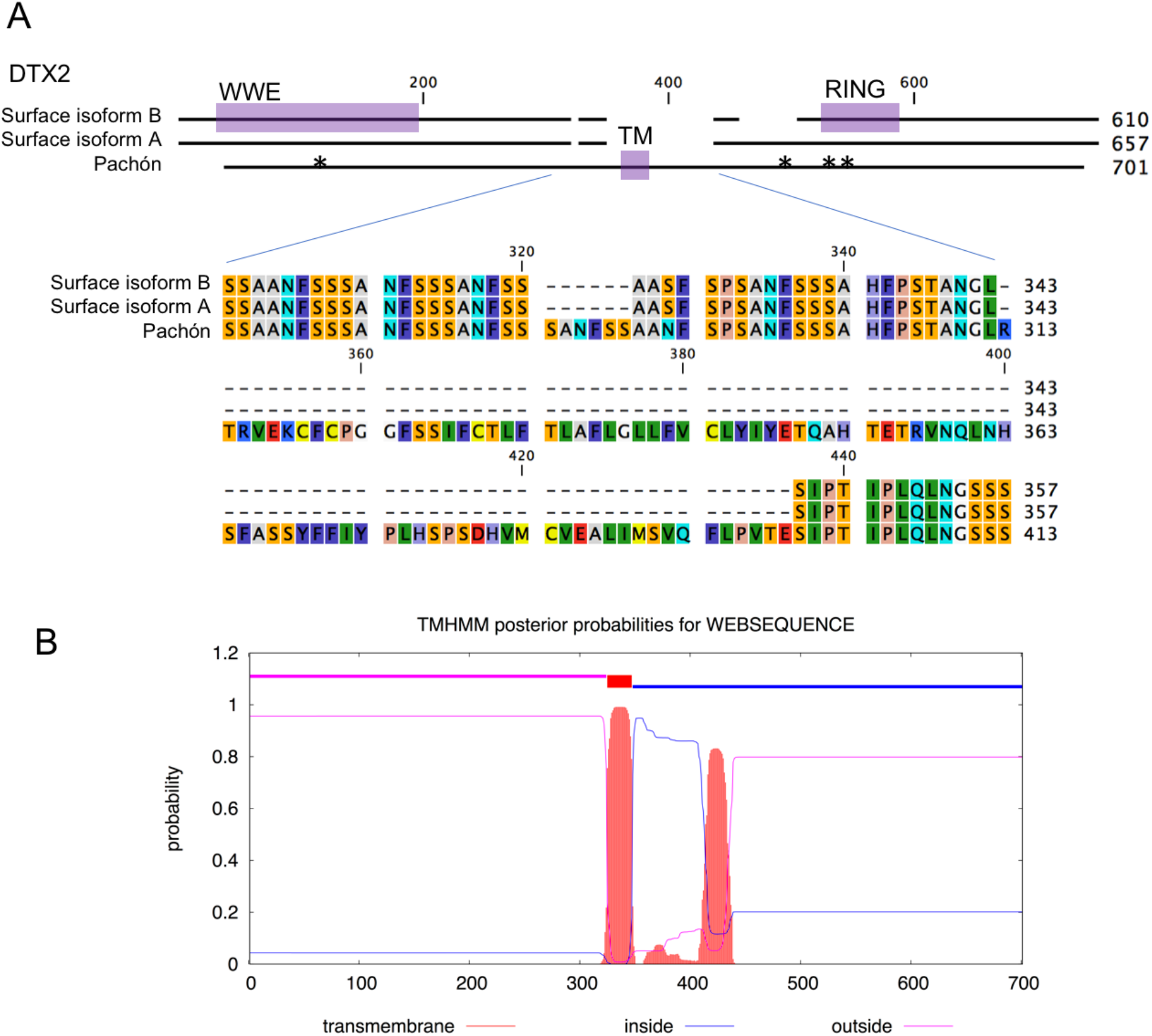
Cavefish have coding changes to DTX2. A, Alignment of predicted protein sequences from surface fish and Pachón cavefish reference genomes. Protein domains are highlighted in purple (WWE, RING, TM: transmembrane). Asterisks indicate position of Pachón amino acid substitutions. B, Results from transmembrane prediction analysis using Pachón cavefish sequence (DTU bioinformatics TMHMM server v. 2.0 http://www.cbs.dtu.dk/services/TMHMM/).

We examined the *dtx2* locus in wild-caught fish from two surface populations (Rio Choy and Rascón), three cavefish populations (Pachón, Tinaja, Molino), a closely related species (*Astyanax aeneus*), and a more distantly related species (*Gymnocorymbus ternetz*, white long fin) using data from Herman et al., 2018 (Table 3, Supplemental data 1). All of the cavefish and one surface fish carry the V116I substitution. We found that only Rio Choy surface have a proline at position 398, suggesting that an alanine at this position may be the ancestral state. We found that the R441H substitution is specific to fish from the Pachón cave and the S443L substitution is only present in Pachón and Tinaja cavefish. We found fish with two, three, and four repeats of the 20bp sequence we identified comparing surface fish and Pachón cavefish. All of the Pachón cavefish have four repeats (n=10).

**Table 3.**
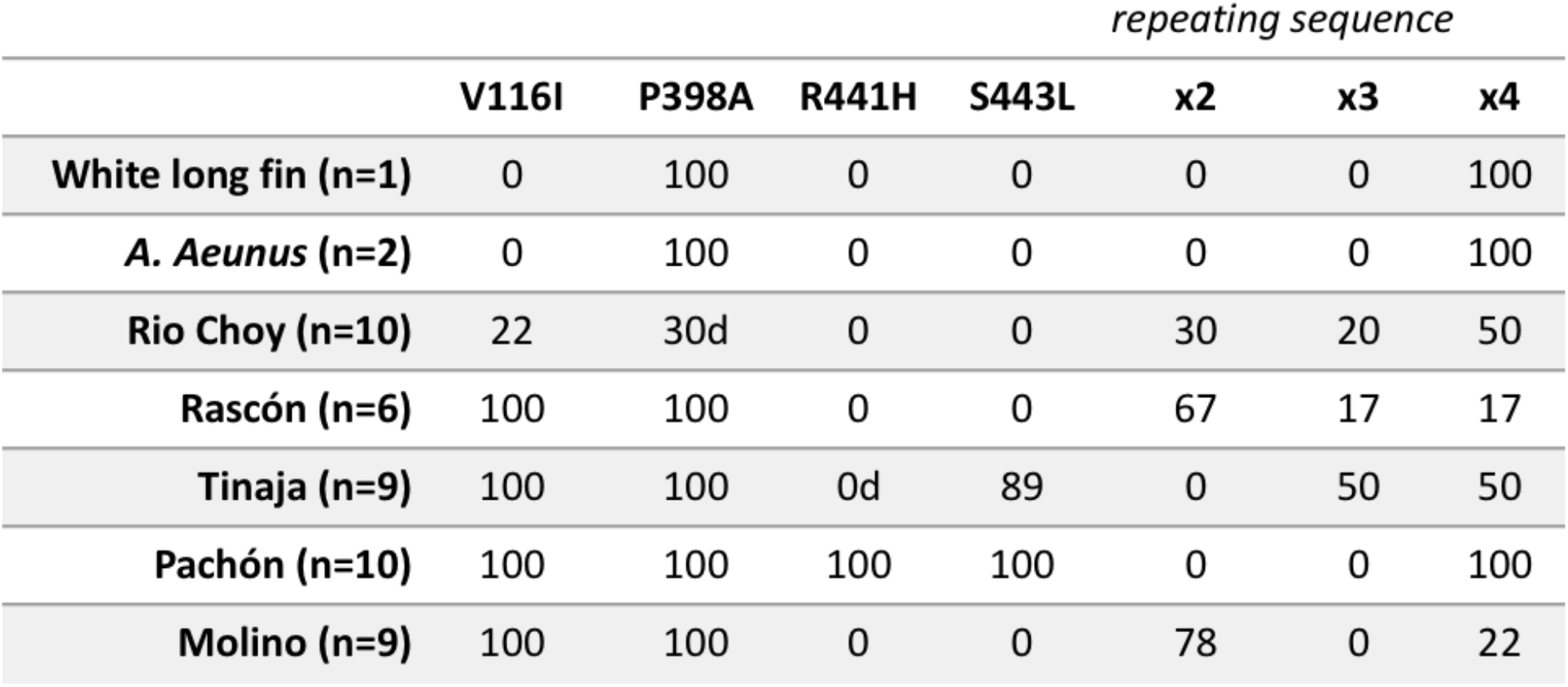
Percentage of fish with the indicated genetic changes in DTX2. “d” indicates that one fish had a deletion at this position.

To search for other genetic changes that could influence carotenoid accumulation in cavefish, we reviewed the literature and generated a list of fifty genes that have known roles in carotenoid processing or are linked with human variation in carotenoid levels (Supplemental table 1) (Bohn et al., 2017; Moran et al., 2018; Toews et al., 2017). None of these genes are on the unplaced scaffolds that were mapped in the fat color QTL. However, two of the genes, *fatty acid elongase 2* (*elovl2*) and *apolipoprotein A-I* (*apoa1a*) are located on chromosome 3 approximately 6mbp and 9mbp from the fat color QTL. We compared the predicted protein sequence of ELOVL2and APOA1A using the available Pachón and surface genomes. There are no coding changes in *apoa1a.* We found that ELOVL2 has two amino acid substitutions (A274V, V281I). However, when we examined the sequences in wild-caught fish we found that all populations have a valine at position 274, including surface populations. Only two Rio Choy surface fish and the outgroups have a valine at position 281. In summary, we found consistent sequence differences in *DTX2* between surface and cave populations but not in *apoa1a* or *elovl2*.

### Expression changes in genes important for carotenoid processing

To more closely examine the molecular basis of carotenoid accumulation, we sought to compare expression of carotenoid-associated genes between surface fish and Tinaja cavefish. We noticed a sharp contrast between the color of the midgut and hindgut in both populations; the hindgut appears more orange suggesting it may be the primary site of carotenoid absorption (Supplemental Figure 2A). We therefore performed RNA sequencing on hindguts from surface, Tinaja, and surface/Tinaja F1 hybrids (n=5 per population). Each sample had over 40 million reads and nearly 90% mapping rate to the *A. mexicanus* genome. We generated a list of normalized gene counts for each sample to control for differences in sequencing depth and used the counts to make comparisons between populations. We detected over 20,000 genes (i.e. at least one read per gene) in each sample. Principal Component Analysis of normalized, variance stabilized data showed that the samples clustered by population (Supplemental Figure 2B).

We performed differential gene expression analysis by first comparing surface fish and Tinaja cavefish samples using DESEq2’s pairwise Wald test. We found that 3373 genes are differentially expressed, 1612 upregulated and 1761 downregulated in cavefish (adjusted p-value <0.05 and log2FoldChange ≥ 0.58). None of the genes within the fat color QTL, including *elovl2* and *apoa1a*, are differentially expressed. We compared expression of genes that are known to regulate carotenoid absorption or that have been associated with carotenoid levels in humans (Supplemental Table 1) and found that five genes are differentially expressed: *ISX*, *scarb1*, *bco1*, *mttp*, and *apoea* (Figure 5). Intestine-specific homeobox gene (*ISX*) is a transcription factor that represses expression *scarb1* and *bco1* (Lobo et al., 2010). SCARB1 is a transmembrane receptor expressed in enterocytes that detects micelles and mediates absorption, and BCO1 cleaves ß-carotene into retinal within enterocytes. We found that expression of *ISX* is significantly higher in Tinaja (1.24-fold) and *scarb1* and *bco1* are significantly lower (−2.63-fold and −2.84-fold) consistent with the regulation of these genes by *ISX*. Tinaja also have reduced expression of two lipid-binding proteins: microsomal triglyceride transfer protein (*mttp*) and apolipoprotein Eb (*apoea*). These proteins function in formation and transport chylomicrons respectively and therefore influence carotenoid transport into the lymphatic system (Figure 5B).

To determine if the expression patterns we observed are influenced by dominant or recessive alleles we used a likelihood ratio test to compare normalized counts between surface, Tinaja, and F1 hybrid samples (Figure 5A). We found that average *ISX* expression level is highest in F1 hybrid samples (normalized counts: 62 CF (cavefish), 125 F1 (hybrid), 18 SF (surface fish), adjusted p-value = .004), but there is considerable variation; two of the samples have levels intermediate between surface and Tinaja and three of the samples are higher. The average normalized counts for the ISX target *scarb1* is intermediate in the hybrid samples (193 CF, 412 F1, 1518 SF, adjusted p-value = 2×10^−11^) and the other target, *bco1* is similar to Tinaja (131 CF, 127 F1, 1282 SF, adjusted p-value = 1×10^−15^). The average normalized counts for the lipid-binding proteins *apoea* (1047 CF, 2081 F1, 2309 SF, adjusted p-value = 0.014) and *mttp* (139 CF, 177 F1, 320 SF, adjusted p-value = 0.006) are intermediate in the F1 hybrid, however two of the F1 samples have normalized *mttp* counts that are higher than the surface fish average. We found an additional gene that was significantly different when we analyzed the samples using the likelihood ratio test: *cd36*, a scavenger receptor that similar to *scarb1* is involved in carotenoid uptake has lower expression in Tinaja and F1 hybrids (4512 Tinaja, 4102 F1, 9354 surface, adjusted p-value =0.03). In summary, we observed that cavefish have altered expression of multiple genes that influence carotenoid absorption, metabolism, and circulation.

## Discussion

When *Astyanax mexicanus* invaded the caves of the Sierra del Abra, they found themselves trapped in a starkly different environment from their ancestral home in the river system. 20-30,000 years later, the populations have evolved strikingly different morphological and metabolic features. A major question is whether these cave traits result from genetic drift, selection, or pleiotropic interactions. Surface adapted *A. mexicanus* utilize carotenoids for vision and pigmentation; two traits that are not under selection in a perpetually dark cave. Although cavefish have reduced or absent melanin pigment, they still exhibit some carotenoid-based pigment. We found that unlike surface fish, Tinaja and Pachón cavefish accumulate carotenoids in the visceral adipose tissue. Here we examined the variables that influence carotenoid-based traits and the genetic and molecular basis of the differences between populations.

Surface-adapted forms of *A. mexicanus* exhibit a bright orange stripe on the anal fin. In a laboratory tank, the most aggressive surface fish develops the most visible stripe, but we found that when fish are housed individually, they each develop a stripe. It is unclear if the amount of food consumed is the only factor in determining stripe formation (i.e. dominant fish obtain the most food), or if the cellular and biochemical pathways involved in behavior or social interaction directly influence carotenoid-processing. It is also unclear the extent to which the stripe itself influences social hierarchy, mate preference or other behaviors.

We observed that Tinaja cavefish fed *ab libitum* accumulate yellow pigment in the visceral adipose tissue. Cave populations experience prolonged nutrient deprivation followed by a sudden influx of food during seasonal floods. This environment has selected for hyperphagia in some populations. Tinaja and Molino cavefish consume more than surface fish in the laboratory and this difference is associated with mutations in melanocortin receptor 4 (MC4R) (Aspiras et al., 2015). MC4R is a G protein-coupled receptor expressed in the hypothalamus that integrates leptin and insulin signals to modulate appetite (Baldini & Phelan, 2019). Human variation at the MC4R locus is associated obesity, anorexia, and one study found a human SNP in MC4R associated with lutein levels in the serum (Borel et al., 2014). Our results show however that differences in MC4R and appetite are not sufficient to explain yellow VAT: Tinaja cavefish VAT remained yellow on a controlled diet, while the Molino cavefish that carry the MC4R mutation, kept on the same diet, have white fat.

Loss of the selective pressures that favor eye formation and function could, in principle, have led to accumulation of mutations or genetic drift in pathways important for production of carotenoid metabolites. Changes to these pathways could, in turn, play a role in eye degeneration. We found that Molino cavefish do not accumulate carotenoids in the VAT showing that eye loss does not necessarily lead to the differences we observed between Surface fish and Tinaja cavefish. We also found no association between eye size and fat color in surface/Tinaja F2 hybrids indicating that the mechanisms that lead to carotenoid accumulation are unlikely to influence eye formation. Eye loss in cavefish is the result of multiple genetic changes, as there have been at least fourteen quantitative trait loci identified that account for variation in eye phenotypes (Protas, Conrad, Gross, Tabin, & Borowsky, 2007). Our current results suggest that carotenoid accumulation is not a pleiotropic phenotype associated with eye loss. However, alterations to retinol production could influence vision. Future studies utilizing hybrid populations may reveal correlations between altered carotenoid processing and visual function.

At early stages of development, cavefish embryos appear more yellow than surface fish embryos. This color difference persists through larval development. As an embryonic trait, the yellow pigment must be passed from the female parent. Consistent with this, cave/surface F1 hybrids that result from a surface fish female are not yellow and those arising from a cavefish female are yellow (see Hinaux et al., 2017). This suggests there is a difference in the carotenoid content in the eggs and provides an opportunity to determine if this has an impact on development by comparing the hybrids produced from cave versus surface females.

Morphological changes to the gut could influence carotenoid accumulation by altering digestion or absorption. For example, Pachón cavefish have altered gut motility at post-larval stages that results in slower transit of food; an adaptation that could increase nutrient absorption (Riddle, Boesmans, Caballero, Kazwiny, & Tabin, 2018). We found that length of the gut is not linked with fat color in surface/Tinaja F2 hybrids, however we identified a candidate gene that could influence development of intestinal cell types: *DTX2*. DTX2 is an E3 ubiquitin ligase that binds to Notch ankyrin repeats via the WWE domain and can mediate Notch activation or degradation depending on the context and interaction with other proteins (Hori et al., 2004; Matsuno et al., 1995; Puca, Chastagner, Meas-Yedid, Israël, & Brou, 2013). Notch signaling in the intestinal epithelium is important for maintenance of crypt-based columnar cells and controls the proportion of secretory and absorptive cell types (Tian et al., 2015). There is evidence that Notch stability and trafficking is controlled by E3 ligases in the intestine but the role of DTX2 specifically has not been investigated (reviewed in Sancho, Cremona, & Behrens, 2015).

We found multiple changes to the DTX2 locus by first comparing the predicted protein sequence between surface fish and Pachón cavefish. We observed additional amino acids in the Pachón sequence that form a predicted transmembrane domain. Numerous transmembrane ubiquitin ligases are present in the mammalian genome that regulate protein degradation and influence cellular processes from membrane trafficking to proliferation (reviewed in Nakamura, 2011). The transmembrane domain in the Pachón cavefish is due to a difference in splicing and consequent changes to the cDNA sequence so it is currently unclear if the transmembrane domain is present in other cavefish populations. We compared the genomic DNA sequence in wild populations to determine if any changes were present in only the Tinaja and Pachón cavefish, and thus correlate with carotenoid accumulation. We found that only Tinaja and Pachón have the S443L substitution present in the RING domain. This residue is conserved between surface fish and humans. The RING domain is important for specifying the activity of the E3 ubiquitin ligase by mediating interaction with E2 ubiquitin conjugating enzymes (Metzger, Pruneda, Klevit, & Weissman, 2014). Future studies will be aimed at understanding how the changes we identified in Pachón influence DTX protein structure, interaction with Notch, and the development of intestinal cell types.

Carotenoid-accumulation could be a pleiotropic consequence of selecting for genetic changes that alter lipid absorption or processing. Carotenoids are absorbed and transported through the same pathways as dietary lipids. Tinaja and Pachón cavefish accumulate fat at earlier stages of development, and have more fat throughout their body as adults compared to surface fish (Aspiras et al., 2015; Hüppop, 2000; Xiong et al., 2018). We found that Molino cavefish do not have yellow VAT and interestingly, they also do not accumulate fat early and have only moderately more fat than surface fish (Xiong et al., 2018). However, counter to the hypothesis that cavefish have increased lipid absorption, we found that Tinaja cavefish have lower expression of the lipid scavengers *scarb1* and *cd36* in the hindgut. We also found reduced expression of microsomal triglyceride transfer protein (*mttp*) and apolipoprotein Eb (*apoea*) that are important for lipid transport. Thus, if Tinaja have increased lipid absorption, they must achieve this through a novel mechanism.

The expression of *scarb1* and *bco1* is inhibited by ISX as a part of a feedback loop that is responsive to changes in carotenoid levels (Lobo et al., 2010). Carotenoids like ß – carotene are absorbed by the action of SCARB1, BCO1 cleaves ß –carotene into retinol and retinol is converted to retinoic acid that binds directly to the promoter of *ISX* to activate expression (Figure 6B). Decreased *bco1* expression is associated with carotenoid accumulation in other species (reviewed in Bohn et al., 2017; Toews et al., 2017). Our results are consistent with the hypothesis that reduced expression of *bco1* contributes to accumulation of carotenoids in cavefish. The regulatory points at which the feedback loop between SCARB1, BCO1, and ISX are altered are currently unclear but it is possible that an increased number of absorptive cells leads to greater carotenoid accumulation and *ISX* expression on the whole tissue level. We found that F1 hybrids have higher expression of ISX than surface fish or cavefish suggesting that there may be multiple antagonistic genes that influence ISX expression. Examining this feedback pathway at other stages of development and under various diets could help dissect how the differences between surface fish and cavefish arise.

**Figure 6.**
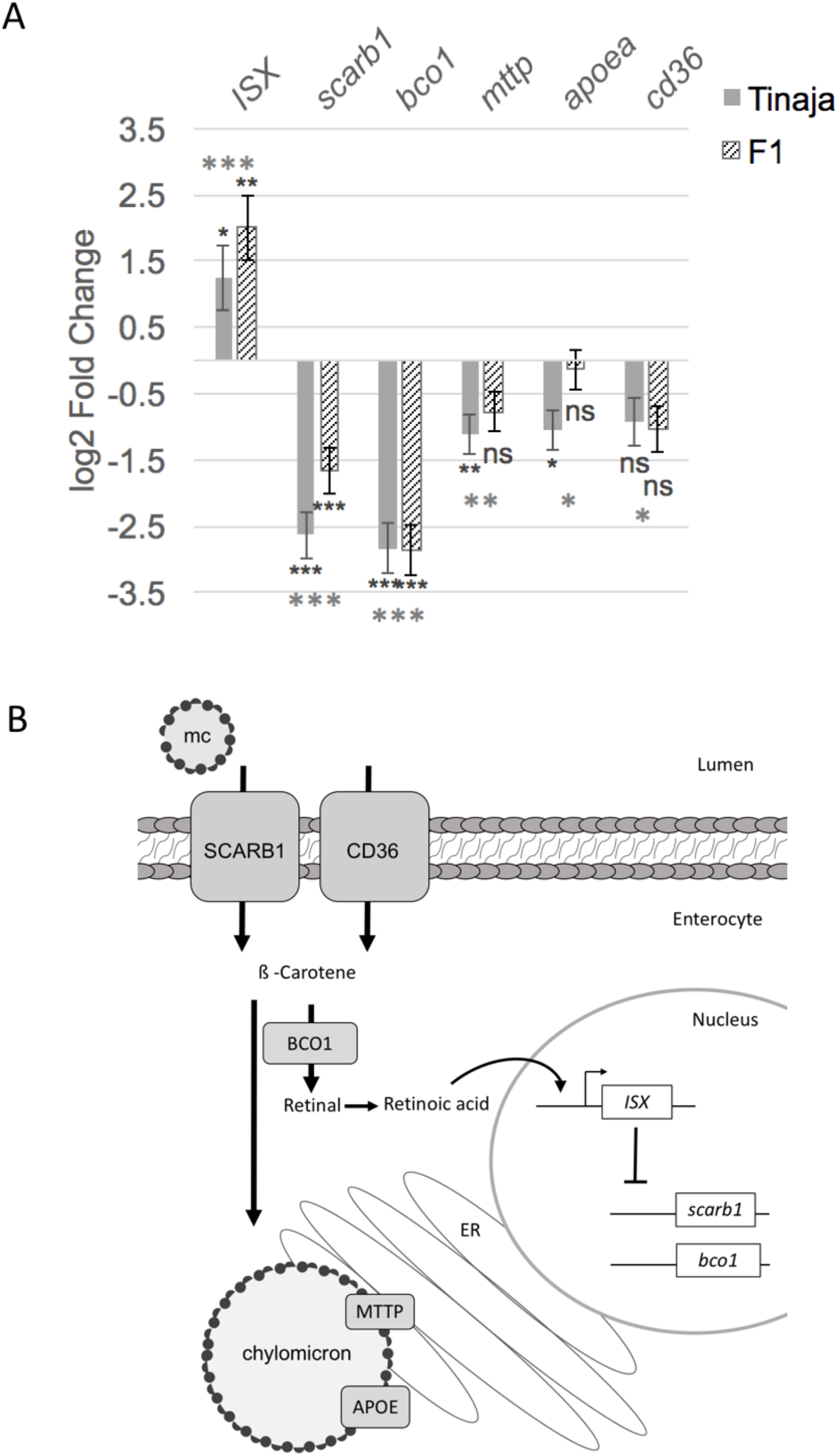
Altered expression of carotenoid-processing genes in Tinaja cavefish. A, Bar graph showing log 2-Fold expression change compared to surface fish for the indicated gene in Tinaja cavefish hindguts and surface/Tinaja F1 hybrid hindguts (n=5 samples per population). Grey asterisks indicate significance from likelihood ratio test comparing all three sample types and black asterisks indicate significance from Wald test comparing to only surface fish (Significance code for adjusted p-value: *<.05 **<.005 ***<.0005, ns>.05). Error bars indicate standard error estimate for log2 fold change. B, Diagram of an enterocyte showing protein functions of the differentially expressed genes. Mixed micelles (mc) containing carotenoids are transported into enterocytes by lipid scavenger proteins SCARB1 and CD36. The carotenoid ß-carotene is cleaved by BCO1 into two molecules of retinal that can form retinoic acid. Retinoic acid binds directly to the ISX promoter to activate expression and ISX protein inhibits expression of *scarb1* and *bco1.* Carotenoids can be packaged into chylomicrons that are formed in the endoplasmic reticulum (ER). MTTP and APOE are proteins associated with chylomicrons that are important for their formation and transport into the lymphatic system.

Finally, could accumulating carotenoids be advantageous in the cave environment? Cavefish have metabolic phenotypes that would be pathological in a human context including: fatty liver (Aspiras et al., 2015), high blood sugar (Riddle, Aspiras, et al., 2018), and increased fat storage (Xiong et al., 2018) yet do not appear to suffer damaging effects. In humans, the health benefits of a diet high in carotenoids are evident but the mechanisms of action are not entirely clear; carotenoids are thought to scavenge free-radicals in tissues and reduce oxidative damage (reviewed in Kaulmann and Bohn, 2014). In the VAT specifically, carotenoids control the transcriptional state of adipocytes and modulate inflammatory pathways (Luisa Bonet, Canas, Ribot, & Palou, 2015). A recent study comparing the immune system of surface-and cave-adapted *A. mexicanus* revealed that Pachón cavefish have fewer pro-inflammatory immune cells in the VAT despite having hypertrophic adipocytes (Peuß et al., 2019). It is intriguing to hypothesize that accumulating carotenoids could provide protective benefits for cavefish. Overall, our findings lay the groundwork for understanding natural variation in carotenoid processing and how it influences metabolism and physiology in a new model system.

## Methods

### Fish husbandry and diet

Fish husbandry was performed according to (Elipot, Legendre, Père, Sohm, & Rétaux, 2014). Unless stated otherwise, fish were fed *ad libitum* with a combination of New Life Spectrum TheraA+ small fish formula and *Artemia* and housed at densities of less than, or equal to, two adult fish per liter of water. For fixed diet experiments fish were housed individually in 1.5L tanks and fed three pellets (approximately 6mg) of New Life Spectrum TheraA+ small fish formula once per day for greater than 4 months. The low carotenoid diet consisted of *Enchytraeus buchholzi* (Fishgobble.com) fed *ad libitum*.

### High Pressure Liquid Chromatography

We extracted pigments from fat samples of the same weight using the following protocol: we added equal parts hexane and methanol to the sample, vortexed the sample for two minute, added an equal volume of water, vortexed the sample for one minute, applied centrifugation at 4,000rpm for 10 minutes, and passed the upper phase through a 0.2-micron nylon filter. The samples were analyzed using an Agilent Technologies 1200 Analytical HPLC with C18 reversed-phase column (4.6mm × 150mm, 80A) with an isocratic mobile phase (acetone:water 50:50 v/v) at a flow rate of 1mL/minute for 45 minutes. UV absorbance was monitored with a MWD at 215nm. We included the following standards dissolved in hexane: HPLC grade retinol (Sigma-Aldrich R7632) and β-carotene (Sigma-Aldrich, 22040).

### Imaging

Images were taken using a Cannon Powershot D12 digital camera. The same illumination and settings were used when comparing images.

#### Quantitative trait loci analysis

##### F2 hybrid population

We bred a surface fish female with a Tinaja cavefish male to produce a clutch of F1 hybrids. We generated a population of surface/Tinaja F2 hybrids by interbreeding individuals from this clutch. The F2 mapping population (n=221) consisted of three clutches produced from breeding paired F1 surface/Tinaja hybrid siblings.

##### Genotype by sequencing

We extracted DNA from caudal tail fins using DNeasy Blood and Tissue DNA extraction kit (Qiagen). DNA was shipped to Novogene (Chula Vista, CA) for quality control analysis and sequencing. Samples that contained greater than 1.5 ug DNA, minimal degradation (as determined by gel electrophoresis), and OD_260_/OD_280_ ratio of 1.8 to 2.0 were used for library construction. Each genomic DNA sample (0.3∼0.6 μg) was digested with Mse1, HaeIII, and EcoR1. Digested fragments were ligated with two barcoded adapters: a compatible sticky end with the primary digestion enzyme and the Illumina P5 or P7 universal sequence. All samples were pooled and size-selected after several rounds of PCR amplification to obtain the required fragments needed to generate DNA libraries. Concentration and insert size of each library was determined using Qubit^®^ 2.0 fluorometer and Agilent^®^ 2100 bioanalyzer respectively. Finally, quantitative real-time PCR (qPCR) was used to detect the effective concentration of each library. Qualified DNA libraries had an effective concentration of greater than 2 nM and were pooled by effective concentration and expected data production. Pair-end sequencing was then performed on Illumina^®^ HiSeq platform, with the read length of 144 bp at each end. Raw Illumina genotype-by-sequencing reads were cleaned and processed through the *process_shortreads* command in the Stacks software package (Catchen, Amores, Hohenlohe, Cresko, & Postlethwait, 2011). The cleaned reads were aligned to the *Astyanax mexicanus* reference genome (AstMex102, INSDC Assembly GCA_000372685.1, Apr 2013) using the Bowtie2 software (Langmead & Salzberg, 2012). The aligned reads of 4 surface fish, 4 Tinaja cavefish and 4 F_1_ surface/Tinaja hybrids were manually stacked using the *pstacks* command. We then assigned the morphotypic origin of each allele and confirmed heterozygosity in the F_1_ samples using the *cstacks* command. Finally, we used this catalog to determine the genotypes at each locus in the F2 samples with the *sstacks* and *genotypes* command. This genotype database was formatted for use in R/qtl (Arends, Prins, Jansen, & Broman, 2010)

##### Linkage map

Using R/qtl, we selected for markers that were homozygous and had opposite genotypes in cavefish versus surface fish (based on three surface and three cave individuals). A linkage map was constructed from these loci using only the F2 population. All markers that were genotyped in less than 180 individuals were omitted, as well as all individuals that had poor marker genotyping (<1500 markers). Markers that did not conform to the expected allele segregation ratio of 1:2:1 were also omitted (p < 1e^−10^). These methods produced a map with 1839 markers, 219 individuals, and 29 linkage groups. Unlinked markers and small linkage groups (<10 markers) were omitted until our map consisted of the optimal 25 linkage groups (*Astyanax mexicanus* has 2n = 50 chromosomes (Kavalco & De Almeida-Toledo, 2007)). Each linkage group had 20-135 markers and markers were reordered by default within the linkage groups, producing a total map length of > 5000 cM. Large gaps in the map were eliminated by manually switching marker order to the best possible order within each linkage group (error.prob = 0.005). The final linkage map consisted of 1800 markers, 219 individuals and 25 linkage groups, spanning 2871 cM. Maximum spacing was 32.1 cM and average spacing was 1.6 cM.

### Sequence analysis of candidate genes

Sequence comparisons were made using Ensembl gene builds for surface fish (Astyanax_mexicanus-2.0, INSDC Assembly <u>GCA_000372685.2</u>, Sep 2017) and Pachón cavefish (AstMex102, INSDC Assembly GCA_000372685.1, Apr 2013). Sequence comparisons between wild-caught fish were made using data from (Herman et al., 2018) and alignments were produced using CLC Sequence Viewer 8.

#### RNA sequencing

##### RNA extraction and cDNA synthesis

Adult *A. mexicanus* were euthanized in 400ppm Tricane and the hindgut was immediately removed and homogenized in 0.3mL Trizol using a motorized pellet pestle and stored at −80°C. Total RNA was extracted using Zymo Research Direct-zol RNA MicroPrep with DNAse treatment according to the manufacturers protocol. LunaScript RT supermix kit with 1μg of RNA was used to synthesize cDNA. All samples were processed on the same day. Diluted cDNA samples (50ng/μL) were used for sequencing.

##### RNA sequencing and differential gene expression analysis

HiSeq Illumina sequencing was performed by Novogene (Chula Vista, CA). All samples were indexed and run as pools, providing an estimated 20-30 million single-end reads per sample.

Samples were processed using an RNA-seq pipeline implemented in the bcbio-nextgen project (https://bcbio-nextgen.readthedocs.org/en/latest/). Raw reads were examined for quality issues using FastQC (http://www.bioinformatics.babraham.ac.uk/projects/fastqc/) to ensure library generation and sequencing are suitable for further analysis.

Reads were aligned to Ensembl build 2 of the *Astyanax mexicanus* genome, augmented with transcript information from Ensembl release 2.0.97 using STAR (Dobin et al., 2013) with soft trimming enabled to remove adapter sequences, other contaminant sequences such as polyA tails and low quality sequences. Alignments were checked for evenness of coverage, rRNA content, genomic context of alignments and complexity using a combination of FastQC, Qualimap (García-Alcalde et al., 2012), MultiQC (https://github.com/ewels/MultiQC), and custom tools. Counts of reads aligning to known genes were generated by featureCounts (Liao, Smyth, & Shi, 2014) and used for further QC with the R package bcbioRNASeq [Steinbaugh MJ, Pantano L, Kirchner RD et al. bcbioRNASeq: R package for bcbio RNA-seq analysis [version 2; peer review: 1 approved, 1 approved with reservations]. F1000Research 2018, 6:1976 (https://doi.org/10.12688/f1000research.12093.2)]. In parallel, Transcripts Per Million (TPM) measurements per isoform were generated by quasialignment using Salmon (Patro, Duggal, & Kingsford, 2015) for downstream differential expression analysis as quantitating at the isoform level has been shown to produce more accurate results at the gene level (Soneson, Love, & Robinson, 2016). Salmon output was imported into R using tximport (Soneson, Love, & Robinson, 2015) and differential gene expression analysis and data visualization was performed using R with the DEseq 2 package (Love, Huber, & Anders, 2014; R-core-team, 2019).

## Acknowledgments

Brian Martineu and Megan Peavey for fish husbandry, Fransisca Leal and Robert Peuß for assisting in fish dissections, and Suzanne McGaugh for sharing sequencing data. Work by JN Hutchinson at the Harvard Chan Bioinformatics Core was funded by the Harvard Medical School Tools and Technology Committee. This work was supported by grants from the National Institutes of Health [HD089934, DK108495].

The authors declare no conflicts of interest.

The data that support the findings of this study are available from the corresponding author upon reasonable request.

---

**Supplemental Figure 1.**
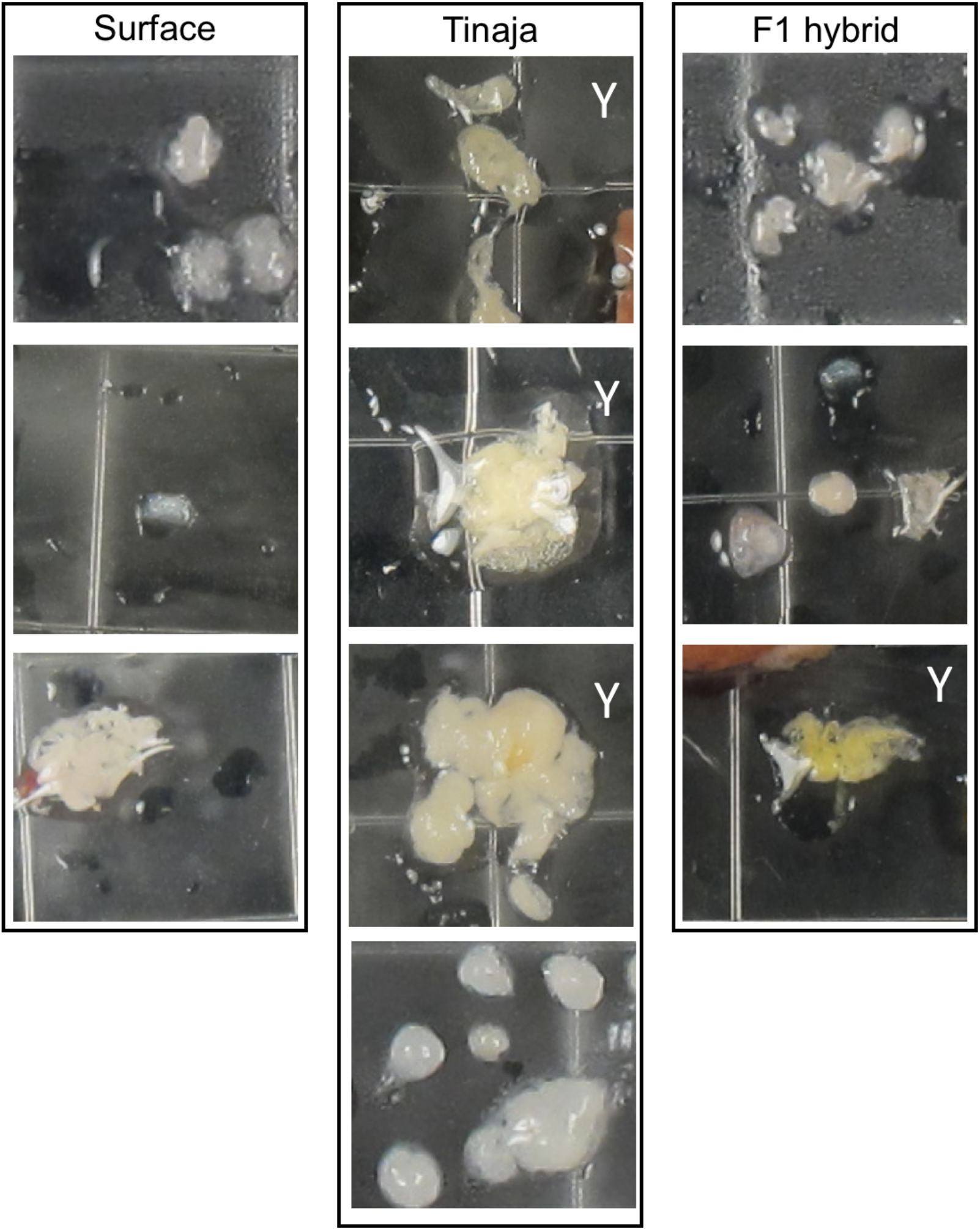
Variation in VAT color among individual surface, Tinaja, and F1 surface/Tinaja hybrids fed a fixed pellet diet. Images are at approximately the same magnification (2.3mm wide). Y indicates which samples are considered yellow.

**Supplemental Figure 2.**
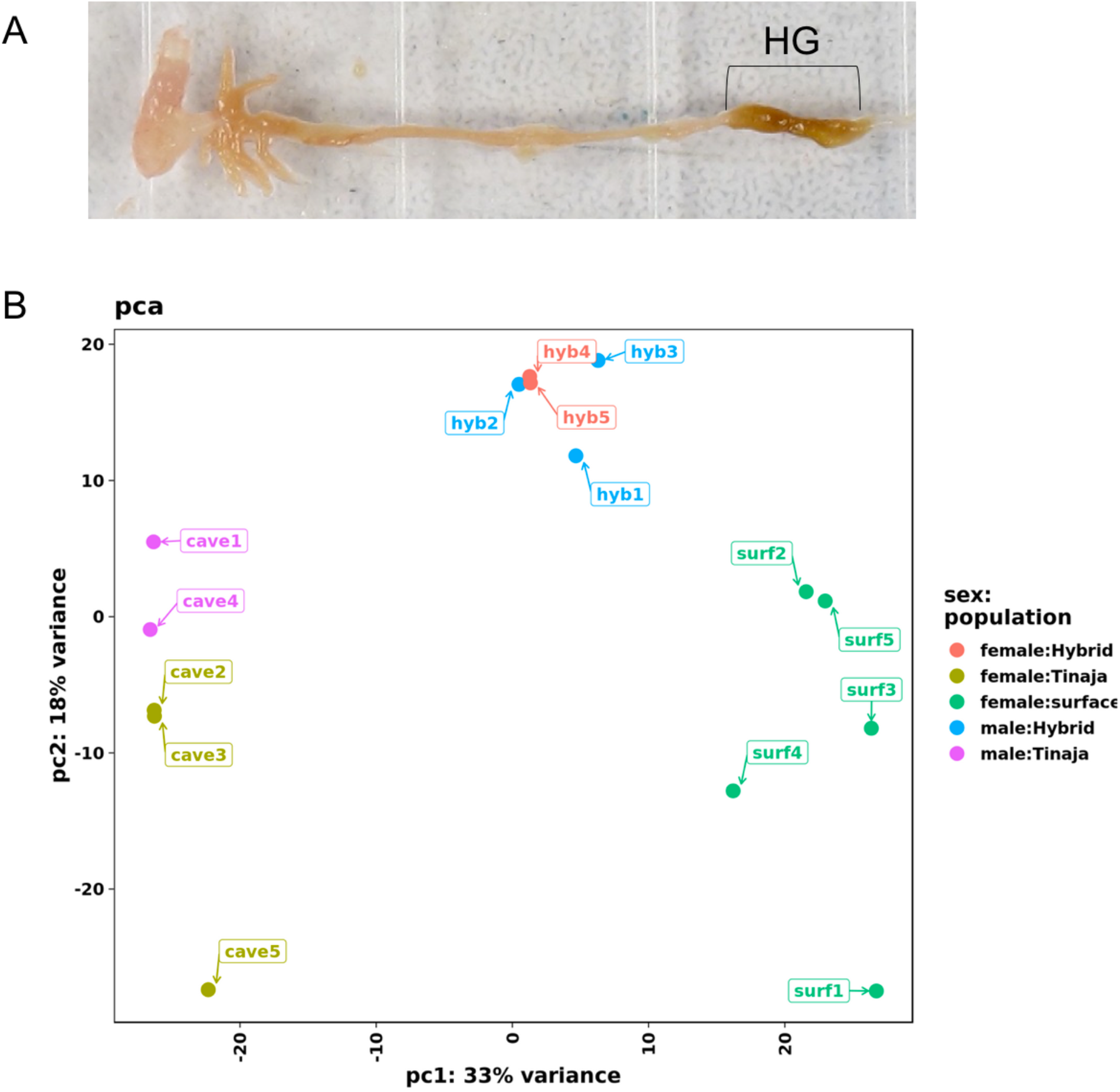
RNA sequencing of hindgut in *A. mexicanus.* A, Image of Tinaja cavefish gastrointestinal tract, HG: hindgut. B, Principle component analysis of normalized, variance stabilized sequencing data from Tinaja (cave), surface (surf), and F1 hybrid (hyb) samples.

## References

Arends, D., Prins, P., Jansen, R. C., & Broman, K. W. (2010). R/qtl: High-throughput multiple QTL mapping. Bioinformatics. https://doi.org/10.1093/bioinformatics/btq565

Aspiras, A. C., Rohner, N., Martineau, B., Borowsky, R. L., & Tabin, C. J. (2015). Melanocortin 4 receptor mutations contribute to the adaptation of cavefish to nutrient-poor conditions. Proceedings of the National Academy of Sciences, 112(31), 9668–9673. https://doi.org/10.1073/pnas.1510802112

Baldini, G., & Phelan, K. D. (2019). The melanocortin pathway and control of appetite-progress and therapeutic implications. Journal of Endocrinology. https://doi.org/10.1530/JOE-18-0596

Berry, S. D., Davis, S. R., Beattie, E. M., Thomas, N. L., Burrett, A. K., Ward, H. E., … Snell, R. G. (2009). Mutation in bovine β-carotene oxygenase 2 affects milk color. Genetics. https://doi.org/10.1534/genetics.109.101741

Bohn, T., Desmarchelier, C., Dragsted, L. O., Nielsen, C. S., Stahl, W., Rühl, R., … Borel, P. (2017). Host-related factors explaining interindividual variability of carotenoid bioavailability and tissue concentrations in humans. Molecular Nutrition and Food Research. https://doi.org/10.1002/mnfr.201600685

Borel, P., Desmarchelier, C., Nowicki, M., Bott, R., Morange, S., & Lesavre, N. (2014). Interindividual variability of lutein bioavailability in healthy men: Characterization, genetic variants involved, and relation with fasting plasma lutein concentration. American Journal of Clinical Nutrition. https://doi.org/10.3945/ajcn.114.085720

Catchen, J. M., Amores, A., Hohenlohe, P., Cresko, W., & Postlethwait, J. H. (2011). Stacks: building and genotyping Loci de novo from short-read sequences. G3 (Bethesda, Md.), 1(3), 171–182. https://doi.org/10.1534/g3.111.000240

Chin, J. S. R., Gassant, C. E., Amaral, P. M., Lloyd, E., Stahl, B. A., Jaggard, J. B., … Duboue, E. R. (2018). Convergence on reduced stress behavior in the Mexican blind cavefish. Developmental Biology, 441(2), 319–327. https://doi.org/10.1016/j.ydbio.2018.05.009

Dobin, A., Davis, C. A., Schlesinger, F., Drenkow, J., Zaleski, C., Jha, S., … Gingeras, T. R. (2013). STAR: Ultrafast universal RNA-seq aligner. Bioinformatics. https://doi.org/10.1093/bioinformatics/bts635

Eggersdorfer, M., & Wyss, A. (2018). Carotenoids in human nutrition and health. Archives of Biochemistry and Biophysics. https://doi.org/10.1016/j.abb.2018.06.001

Elipot, Y., Legendre, L., Père, S., Sohm, F., & Rétaux, S. (2014). *Astyanax* Transgenesis and Husbandry: How Cavefish Enters the Laboratory. Zebrafish, 11(4), 291–299. https://doi.org/10.1089/zeb.2014.1005

Elliott, W. R. (2015). Cave Biodiversity and Ecology of the Sierra de El Abra Region. In Biology and Evolution of the Mexican Cavefish (pp. 59–76). https://doi.org/10.1016/B978-0-12-802148-4.00003-7

Eriksson, J., Larson, G., Gunnarsson, U., Bed’hom, B., Tixier-Boichard, M., Strömstedt, L., … Andersson, L. (2008). Identification of the Yellow skin gene reveals a hybrid origin of the domestic chicken. PLoS Genetics. https://doi.org/10.1371/journal.pgen.1000010

García-Alcalde, F., Okonechnikov, K., Carbonell, J., Cruz, L. M., Götz, S., Tarazona, S., … Conesa, A. (2012). Qualimap: Evaluating next-generation sequencing alignment data. Bioinformatics. https://doi.org/10.1093/bioinformatics/bts503

Gore, A. V, Tomins, K. A., Iben, J., Ma, L., Castranova, D., Davis, A. E., … Weinstein, B. M. (2018). An epigenetic mechanism for cavefish eye degeneration. Nature Ecology & Evolution, 2(7), 1155–1160. https://doi.org/10.1038/s41559-018-0569-4

Gross, J. B., & Powers, A. K. (2018). A Natural Animal Model System of Craniofacial Anomalies: The Blind Mexican Cavefish. Anatomical Record. https://doi.org/10.1002/ar.23998

GTEx Analysis Release V8 (dbGaP Accession phs000424.v8.p2). (n.d.).

Herman, A., Brandvain, Y., Weagley, J., Jeffery, W. R., Keene, A. C., Kono, T. J. Y., … McGaugh, S. E. (2018). The role of gene flow in rapid and repeated evolution of cave-related traits in Mexican tetra, Astyanax mexicanus. Molecular Ecology, 27(22), 4397–4416. https://doi.org/10.1111/mec.14877

Hinaux, H., Pottin, K., Chalhoub, H., Père, S., Elipot, Y., Legendre, L., & Rétaux, S. (2011). A Developmental Staging Table for *Astyanax mexicanus* Surface Fish and Pachón Cavefish. Zebrafish, 8(4), 155–165. https://doi.org/10.1089/zeb.2011.0713

Hinaux, H., Recher, G., Alié, A., Legendre, L., Blin, M., & Rétaux, S. (2017). Lens apoptosis in the Astyanax blind cavefish is not triggered by its small size or defects in morphogenesis. PLoS ONE. https://doi.org/10.1371/journal.pone.0172302

Hori, K., Fostier, M., Ito, M., Fuwa, T. J., Go, M. J., Okano, H., … Matsuno, K. (2004). Drosophila deltex mediates suppressor of hairless-independent and late-endosomal activation of notch signaling. Development. https://doi.org/10.1242/dev.01448

Hüppop, K. (2000). How do cave animals cope with the food scarcity in caves? In Ecosystems of the World: Subterranean ecosystems.

Hyacinthe, C., Attia, J., & Rétaux, S. (2019). Evolution of acoustic communication in blind cavefish. Nature Communications. https://doi.org/10.1038/s41467-019-12078-9

Jeffery, W. R., Ma, L., Parkhurst, A., & Bilandžija, H. (2016). Pigment Regression and Albinism in Astyanax Cavefish. In Biology and Evolution of the Mexican Cavefish (pp. 155–173). Elsevier. https://doi.org/10.1016/B978-0-12-802148-4.00008-6

Kaulmann, A., & Bohn, T. (2014). Carotenoids, inflammation, and oxidative stress-implications of cellular signaling pathways and relation to chronic disease prevention. Nutrition Research. https://doi.org/10.1016/j.nutres.2014.07.010

Kavalco, K. F., & De Almeida-Toledo, L. F. (2007). Molecular cytogenetics of blind Mexican tetra and comments on the karyotypic characteristics of genus Astyanax (Teleostei, Characidae). Zebrafish. https://doi.org/10.1089/zeb.2007.0504

Klaassen, H., Wang, Y., Adamski, K., Rohner, N., & Kowalko, J. E. (2018). CRISPR mutagenesis confirms the role of oca2 in melanin pigmentation in Astyanax mexicanus. Developmental Biology. https://doi.org/10.1016/j.ydbio.2018.03.014

Krishnan, J., & Rohner, N. (2017). Cavefish and the basis for eye loss. Philosophical Transactions of the Royal Society B: Biological Sciences. https://doi.org/10.1098/rstb.2015.0487

Langmead, B., & Salzberg, S. L. (2012). Fast gapped-read alignment with Bowtie 2. Nature Methods, 9(4), 357–359. https://doi.org/10.1038/nmeth.1923

Liao, Y., Smyth, G. K., & Shi, W. (2014). FeatureCounts: An efficient general purpose program for assigning sequence reads to genomic features. Bioinformatics. https://doi.org/10.1093/bioinformatics/btt656

Lloyd, E., Olive, C., Stahl, B. A., Jaggard, J. B., Amaral, P., Duboué, E. R., & Keene, A. C. (2018). Evolutionary shift towards lateral line dependent prey capture behavior in the blind Mexican cavefish. Developmental Biology, 441(2), 328–337. https://doi.org/10.1016/j.ydbio.2018.04.027

Lobo, G. P., Hessel, S., Eichinger, A., Noy, N., Moise, A. R., Wyss, A., … Von Lintig, J. (2010). ISX is a retinoic acid-sensitive gatekeeper that controls intestinal β,β-carotene absorption and vitamin A production. FASEB Journal. https://doi.org/10.1096/fj.09-150995

Love, M. I., Huber, W., & Anders, S. (2014). Moderated estimation of fold change and dispersion for RNA-seq data with DESeq2. Genome Biology. https://doi.org/10.1186/s13059-014-0550-8

Luisa Bonet, M., Canas, J. A., Ribot, J., & Palou, A. (2015). Carotenoids and their conversion products in the control of adipocyte function, adiposity and obesity. Archives of Biochemistry and Biophysics. https://doi.org/10.1016/j.abb.2015.02.022

Lyon, A., Powers, A. K., Gross, J. B., & O’Quin, K. E. (2017). Two - Three loci control scleral ossicle formation via epistasis in the cavefish astyanax mexicanus. PLoS ONE, 12(2). https://doi.org/10.1371/journal.pone.0171061

Matsuno, K., Diederich, R. J., Go, M. J., Blaumueller, C. M., & Artavanis-Tsakonas, S. (1995). Deltex acts as a positive regulator of Notch signaling through interactions with the Notch ankyrin repeats. Development.

Metzger, M. B., Pruneda, J. N., Klevit, R. E., & Weissman, A. M. (2014). RING-type E3 ligases: Master manipulators of E2 ubiquitin-conjugating enzymes and ubiquitination. Biochimica et Biophysica Acta - Molecular Cell Research. https://doi.org/10.1016/j.bbamcr.2013.05.026

Moran, N. E., Mohn, E. S., Hason, N., Erdman, J. W., & Johnson, E. J. (2018). Intrinsic and extrinsic factors impacting absorption, metabolism, and health effects of dietary carotenoids. Advances in Nutrition. https://doi.org/10.1093/ADVANCES/NMY025

Nakamura, N. (2011). The role of the transmembrane RING finger proteins in cellular and organelle function. Membranes. https://doi.org/10.3390/membranes1040354

Patro, R., Duggal, G., & Kingsford, C. (2015). Accurate, fast, and model-aware transcript expression quantification with Salmon. BioRxiv. https://doi.org/10.1101/021592

Peuß, R., Box, A. C., Wang, Y., Chen, S., Krishnan, J., Tsuchiya, D., … Rohner, N. (2019). Single cell analysis reveals modified hematopoietic cell composition affecting inflammatory and immunopathological responses in *Astyanax mexicanus*. BioRxiv, 647255. https://doi.org/10.1101/647255

Protas, M., Conrad, M., Gross, J. B., Tabin, C., & Borowsky, R. (2007). Regressive Evolution in the Mexican Cave Tetra, Astyanax mexicanus. Current Biology, 17(5), 452–454. https://doi.org/10.1016/j.cub.2007.01.051

Puca, L., Chastagner, P., Meas-Yedid, V., Israël, A., & Brou, C. (2013). α-arrestin 1 (ARRDC1) and β-arrestins cooperate to mediate Notch degradation in mammals. Journal of Cell Science. https://doi.org/10.1242/jcs.130500

R-core-team. (2019). R: A Language and Environment for Statistical Computing. Vienna, Austria: R Foundation for Statistical Computing. Retrieved from https://www.r-project.org/

Riddle, M. R., Aspiras, A. C., Gaudenz, K., Peuß, R., Sung, J. Y., Martineau, B., … Rohner, N. (2018). Insulin resistance in cavefish as an adaptation to a nutrient-limited environment, Nature, 555(7698), 647–651. https://doi.org/10.1038/nature26136

Riddle, M. R., Boesmans, W., Caballero, O., Kazwiny, Y., & Tabin, C. J. (2018). Morphogenesis and motility of the Astyanax mexicanus gastrointestinal tract. Developmental Biology, 441(2), 285–296. https://doi.org/10.1016/j.ydbio.2018.06.004

Sakudoh, T., Kuwazaki, S., Iizuka, T., Narukawa, J., Yamamoto, K., Uchino, K., … Tsuchida, K. (2013). CD36 homolog divergence is responsible for the selectivity of carotenoid species migration to the silk gland of the silkworm Bombyx mori. Journal of Lipid Research. https://doi.org/10.1194/jlr.M032771

Sanchez-Pulido, L., & Ponting, C. P. (2018). TMEM132: An ancient architecture of cohesin and immunoglobulin domains define a new family of neural adhesion molecules. Bioinformatics. https://doi.org/10.1093/bioinformatics/btx689

Sancho, R., Cremona, C. A., & Behrens, A. (2015). Stem cell and progenitor fate in the mammalian intestine: Notch and lateral inhibition in homeostasis and disease. EMBO Reports. https://doi.org/10.15252/embr.201540188

Sefc, K. M., Brown, A. C., & Clotfelter, E. D. (2014). Carotenoid-based coloration in cichlid fishes. Comparative Biochemistry and Physiology Part A: Molecular & Integrative Physiology, 173, 42–51. https://doi.org/10.1016/j.cbpa.2014.03.006

Smith, C. M., Hayamizu, T. F., Finger, J. H., Bello, S. M., McCright, I. J., Xu, J., … Ringwald, M. (2019). The mouse Gene Expression Database (GXD): 2019 update. Nucleic Acids Research. https://doi.org/10.1093/nar/gky922

Soneson, C., Love, M. I., & Robinson, M. D. (2015). Differential analyses for RNA-seq: transcript-level estimates improve gene-level inferences. F1000Research. https://doi.org/10.12688/f1000research.7563.1

Soneson, C., Love, M. I., & Robinson, M. D. (2016). Differential analyses for RNA-seq: Transcript-level estimates improve gene-level inferences [version 2; referees: 2 approved]. F1000Research. https://doi.org/10.12688/F1000RESEARCH.7563.2

Sonnhammer, E. L., von Heijne, G., & Krogh, A. (1998). A hidden Markov model for predicting transmembrane helices in protein sequences. Proceedings / … International Conference on Intelligent Systems for Molecular Biology; ISMB. International Conference on Intelligent Systems for Molecular Biology.

Stahl, B. A., & Gross, J. B. (2017). A Comparative Transcriptomic Analysis of Development in Two Astyanax Cavefish Populations. Journal of Experimental Zoology Part B: Molecular and Developmental Evolution. https://doi.org/10.1002/jez.b.22749

Stahl, B. A., Jaggard, J. B., Chin, J. S. R., Kowalko, J. E., Keene, A. C., & Duboué, E. R. (2019). Manipulation of Gene Function in Mexican Cavefish. Journal of Visualized Experiments, (146). https://doi.org/10.3791/59093

Sundvold, H., Helgeland, H., Baranski, M., Omholt, S. W., & Våge, D. I. (2011). Characterisation of a novel paralog of scavenger receptor class B member I (SCARB1) in Atlantic salmon (Salmo salar). BMC Genetics. https://doi.org/10.1186/1471-2156-12-52

Tian, H., Biehs, B., Chiu, C., Siebel, C. W., Wu, Y., Costa, M., … Klein, O. D. (2015). Opposing activities of notch and wnt signaling regulate intestinal stem cells and gut homeostasis. Cell Reports. https://doi.org/10.1016/j.celrep.2015.03.007

Toews, D. P. L., Hofmeister, N. R., & Taylor, S. A. (2017). The Evolution and Genetics of Carotenoid Processing in Animals. Trends in Genetics. https://doi.org/10.1016/j.tig.2017.01.002

Turner, B. J. (2017). Biology and Evolution of the Mexican Cavefish. Edited by Alex C. Keene, Masato Yoshizawa, and Suzanne E. McGaugh. Academic Press. Amsterdam (The Netherlands) and Boston (Massachusetts): Elsevier. $99.95. xiv + 403 p.; ill.; index. ISBN: 978-0-12-802148-. The Quarterly Review of Biology, 92(4), 485–486. https://doi.org/10.1086/694993

Våge, D. I., & Boman, I. A. (2010). A nonsense mutation in the beta-carotene oxygenase 2 (BCO2) gene is tightly associated with accumulation of carotenoids in adipose tissue in sheep (Ovis aries). BMC Genetics. https://doi.org/10.1186/1471-2156-11-10

Vandeweyer, G., Van Der Aa, N., Reyniers, E., & Kooy, R. F. (2012). The contribution of CLIP2 haploinsufficiency to the clinical manifestations of the Williams-Beuren syndrome. American Journal of Human Genetics. https://doi.org/10.1016/j.ajhg.2012.04.020

Wolfe, D. A., & Cornwell, D. G. (1964). Carotenoids of Cavernicolous Crayfish. Science, 144(3625), 1467–1469. https://doi.org/10.1126/science.144.3625.1467

Xiong, S., Krishnan, J., Peuß, R., & Rohner, N. (2018). Early adipogenesis contributes to excess fat accumulation in cave populations of Astyanax mexicanus. Developmental Biology, 441(2), 297–304. https://doi.org/10.1016/j.ydbio.2018.06.003

Yong, L., Peichel, C. L., & McKinnon, J. S. (2016). Genetic architecture of conspicuous red ornaments in female threespine stickleback. G3: Genes, Genomes, Genetics. https://doi.org/10.1534/g3.115.024505

Yoshizawa, M., Robinson, B. G., Duboué, E. R., Masek, P., Jaggard, J. B., O’Quin, K. E., … Keene, A. C. (2015). Distinct genetic architecture underlies the emergence of sleep loss and prey-seeking behavior in the Mexican cavefish. BMC Biology, 13, 15. https://doi.org/10.1186/s12915-015-0119-3

